# *Wolbachia* control stem cell behavior and stimulate germline proliferation in filarial nematodes

**DOI:** 10.1101/244046

**Authors:** Foray Vincent, Pérez-Jiménez Mercedes M., Fattouh Nour, Landmann Frédéric

## Abstract

Although symbiotic interactions are ubiquitous in the living world, examples of developmental symbioses are still scarce. We show here the crucial role of *Wolbachia* in the oogenesis of filarial nematodes, a class of parasites of biomedical and veterinary relevance. While the *Wolbachia*-depleted nematodes produce faulty embryos, we identified thanks to newly generated techniques the earliest requirements of *Wolbachia* in the germline. They stimulate its proliferation in a cell-autonomous manner, in parallel of the known key controllers, and not through nucleotide supplementation as previously hypothesized. We also found *Wolbachia* to maintain the quiescence of a pool of germline stem cells ensuring for many years a constant delivery of about 1400 eggs per day. The loss of quiescence upon *Wolbachia* depletion, as well as the disorganization of the distal germline suggest that *Wolbachia* are required to execute the proper germline stem cell developmental program in order to produce viable eggs and embryos.

## INTRODUCTION

Symbiotic interactions between metazoans and prokaryotes have shaped the living world (Hurst, 2017; McFall-Ngai, 2015). Considered for a long time of a too overwhelming complexity to be studied at cellular and molecular levels, the study of symbiosis remained confined to the field of ecology, until developmental and evolutionary biologists took advantage of the available genomes and the "omics" revolution (McFall-Ngai et al., 2013). Bacterial and animal communities have coevolved to sometimes reach an intimate interdependency. Whether acquired from the environment or vertically transmitted through the female germline, symbionts have established a wide continuum of interactions with their hosts, from parasitism to mutualism. Understanding the mechanisms underlying these stable associations, and the benefits for each partners often appear challenging, but efforts are nonetheless redoubled when economical or biomedical interests are at stake (Slatko et al., 2014). Such is the case of symbioses involving *Wolbachia*, a genus of Gram-negative alphaproteobacteria present in a plethora of invertebrate hosts. Up to date the ever-growing number of *Wolbachia* strains has been classified into 14 supergroups by MLST sequencing, reflecting their diversity across taxa (Baldo et al., 2006). In terrestrial arthropods species, these facultative endosymbionts evolved sex-ratio distortion strategies to favor their vertical transmission through the female progeny (Werren et al., 2008). The discovery that *Wolbachia* from *D. melanogaster* newly introduced into mosquito vectors are able to interfere with the transmission of arboviruses contributed to the popularity of these bacteria (Kamtchum-Tatuene et al., 2016). Aside from these facultative interactions, the coevolution of *Wolbachia* with their host and their transovarian mode of transmission have led to developmental symbioses, where the symbionts have become necessary for the making of an egg. This was reported for the first time in *Asobara tabida*, a parasitic wasp relying on *Wolbachia* to achieve its oogenesis otherwise apoptotic (Dedeine et al., 2001). While a plethora of developmental symbioses probably awaits to be discovered, current evidences are still scarce. The *Vibrio fisheri*-squid symbiosis remains the most achieved comprehension of such interactions. The free-living bacteria harvested by the juvenile cephalopod allow the development the light organ they colonize, and both partners undergo developmental and transcriptional changes in response to the symbiosis (McFall-Ngai, 2014).

Because *Wolbachia* are still genetically intractable, most of the cell biology of the interaction with the host we know as well as our knowledge of their intracellular lifestyle derive mostly from studies either in the fruit fly or in insect cell cultures (Ferree et al., 2005; Geoghegan et al., 2017). Very few *Wolbachia* effectors have been characterized to date, and restricted to interactions with insects (Beckmann et al., 2017; Lepage et al., 2017; Ote et al., 2016). Sometimes the complex mode of life of the host itself is an impediment to explore the basis of a symbiosis. This explains why our understanding of the filarial nematode-*Wolbachia* interaction at the cell level is still in its infancy, although discovered 40 years ago (Kozek, 1977). These species of parasitic nematodes belong to the Onchocercidae family. Transmitted by blood feeding arthropods to a vertebrate host where they develop and live as adults in the lymph or the blood, they cause filariasis, the most debilitating tropical neglected diseases in humans, and a burden in animals, fatal in dogs and cats (McCall et al., 2008; Slatko et al., 2010). Genomic and phylogenetic studies support the occurence of several independent associations of Onchocercidae with *Wolbachia*, followed by secondary losses, leading to current estimates of 37% of filarial species living in symbiosis with *Wolbachia* (Ferri et al., 2011). Yet all species infecting humans but one harbor *Wolbachia*. Antibiotic therapies revealed a mutualistic relationship between *Wolbachia* and filarial nematodes, since worms depleted of their symbionts have been described as sterile due to apoptosis in embryos, and present a longevity reduced from about 10 years to few months (Landmann et al., 2011; Taylor et al., 2010). This highlights a requirement of *Wolbachia* to allow embryonic development and to support the adult survival, consistent with their localization in the female germline and the somatic hypodermal chords. In absence of *Wolbachia*, apoptosis is the ultimate fate of developing embryos and worms stop releasing viable microfilariae, however females still produce embryos with a fraction of them displaying polarity defect phenotypes at early stages suggesting possible earlier requirements of the endosymbionts, i.e. during oogenesis (Landmann et al., 2014). Because of the transovarian transmission of *Wolbachia*, their abundance and tropism for the germline (Fischer et al., 2011; Landmann et al., 2010), we hypothesized they may regulate some fundamental aspects of the germline stem cell behavior and may subsequently affect the proliferation and the production of eggs in filarial species.

We chose to explore this developmental symbiosis using *Brugia malayi*, a causative agent of human filariasis, because it is the sole filarial parasite infecting humans that can be raised in the laboratory with a surrogate rodent host. Both the *Wolbachia* and the worm genomes are sequenced and annotated, and artificial aposymbiotic worms can be obtained by antibiotic treatment under laboratory conditions (Foster et al., 2005; Ghedin et al., 2007). The tetracycline regimen used to deplete *Wolbachia* has been shown to have no direct effect on the fertility of the worm when applied to the naturally *Wolbachia-free A. viteae* filarial species (Hoerauf et al., 1999).

We developed new techniques of dissection, permeabilization and staining for these large and unwieldy nematodes, as well as imaging processing programs in order to characterize precisely their fruitful gametogenesis in whole-mount gonads, otherwise previously observed in cross-sections only. This allowed us to establish that ovaries contain a distal pool of hundreds of quiescent germline stem cells, which once activated proliferate while in transit along a mitotic proliferative zone made of about 3,000 nuclei, until commitment into the meiotic program. We found the distal tip cell to be dispensable to the germ cell proliferation, a marked evolutionary difference with the nematode *C. elegans*. Nonetheless the proliferation remained under the control of a Notch signaling pathway, possibly emanating from the distal somatic gonad, and under the control of *Wolbachia* in a cell-autonomous manner. The depletion of *Wolbachia* caused not only a reduced proliferation, but alleviated the germline stem cell quiescence, and disturbed the germline organization without triggering apoptosis, suggesting that the endosymbionts have become essential to execute the correct germ stem cell developmental program leading otherwise to a decreased production of potentially faulty oocytes.

## RESULTS

### Depletion of *Wolbachia* leads to a decrease of germline cell proliferation

*B. malayi* females possess two ovaries, and their distal parts are located in the posterior of the body. Males however display a single testis whose distal end lies in the anterior of the body (Figure 1A). Both females and males harbor *Wolbachia* in lateral thickenings of the somatic hypodermal tissues -the hypodermal chords-. While the endosymbionts are absent from the male testis, they occupy the female germline, with the highest titer in the distal ovaries (Figures 1B-1D, 38 bacteria/germline nucleus in average along the most distal 100µm). To explore the contribution of *Wolbachia* to the fertility of their host and their impact on the gametogenesis, we compared gonads from *Wolbachia*-depleted adult worms to their wild-type counterparts. The depletion is obtained by applying a standard protocol consisting in a 6-week tetracycline treatment followed by a 2-week clearance period in order to avoid any direct effect of the antibiotics (Figure S1). Instead of classical microscopy observations on body cross sections obtained from paraffin-embedded worms, we developed a new set of techniques ultimately allowing us to access cell dynamics parameters during germline development. To assess the effect of *Wolbachia* depletion on the germline proliferation, we first stained ovaries with an anti-phospho histone 3 (PH3) antibody to reveal mitotic nuclei, and we imaged the distal ovary tips containing the PH3-positive nuclei (Figure 2B). We found an average loss of 32.5 % of PH3+ nuclei from wild-type to *Wolbachia-depleted* – *Wb*(*-*) – ovaries (Figure 2C). In order to elucidate the origin of a decrease in mitotic events, and gain insights into the defects caused by the *Wolbachia* depletion, we characterized the germline development in *B. malayi*.

**Figure 1.**
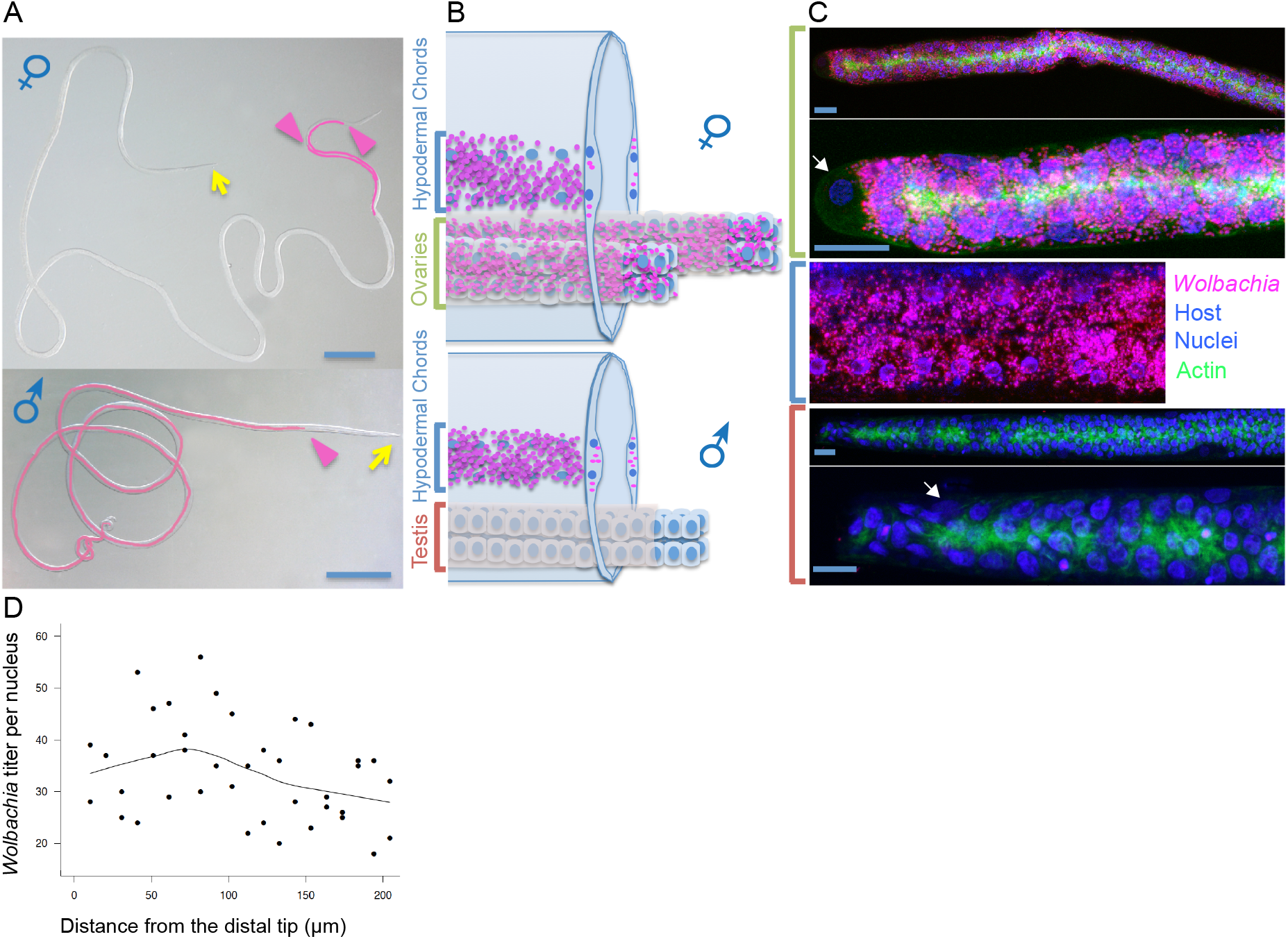
*Wolbachia* are present in the hypodermal chords of both *B. malayi* male and female worms but are restricted to the female germline. (A) Localization of the female ovaries and the male reproductive system in pink on bright field micrographs of worms, with arrowheads indicated the distal part of the gonads. Yellow arrows point to the anterior part of the worms. (B) Schemes showing cross-sections of adult worms and *Wolbachia* -magenta foci-distribution. (C) Confocal images of an ovary distal tip (top), a hypodermal chord -middle-, and a testis distal tip -bottom-, revealing *Wolbachia* - magenta-, host nuclei -blue- and actin -green, see material and methods-. White arrows point to the Distal Tip Cell -DTC-, acting as a germline stem cell niche in nematodes. Scale bars = 1mm (A) and 10 μm (C). (D) *Wolbachia* density in the distal ovary, observed as propidium iodide foci per host nuclei as a function of length expressed in micrometers.

**Figure 2.**
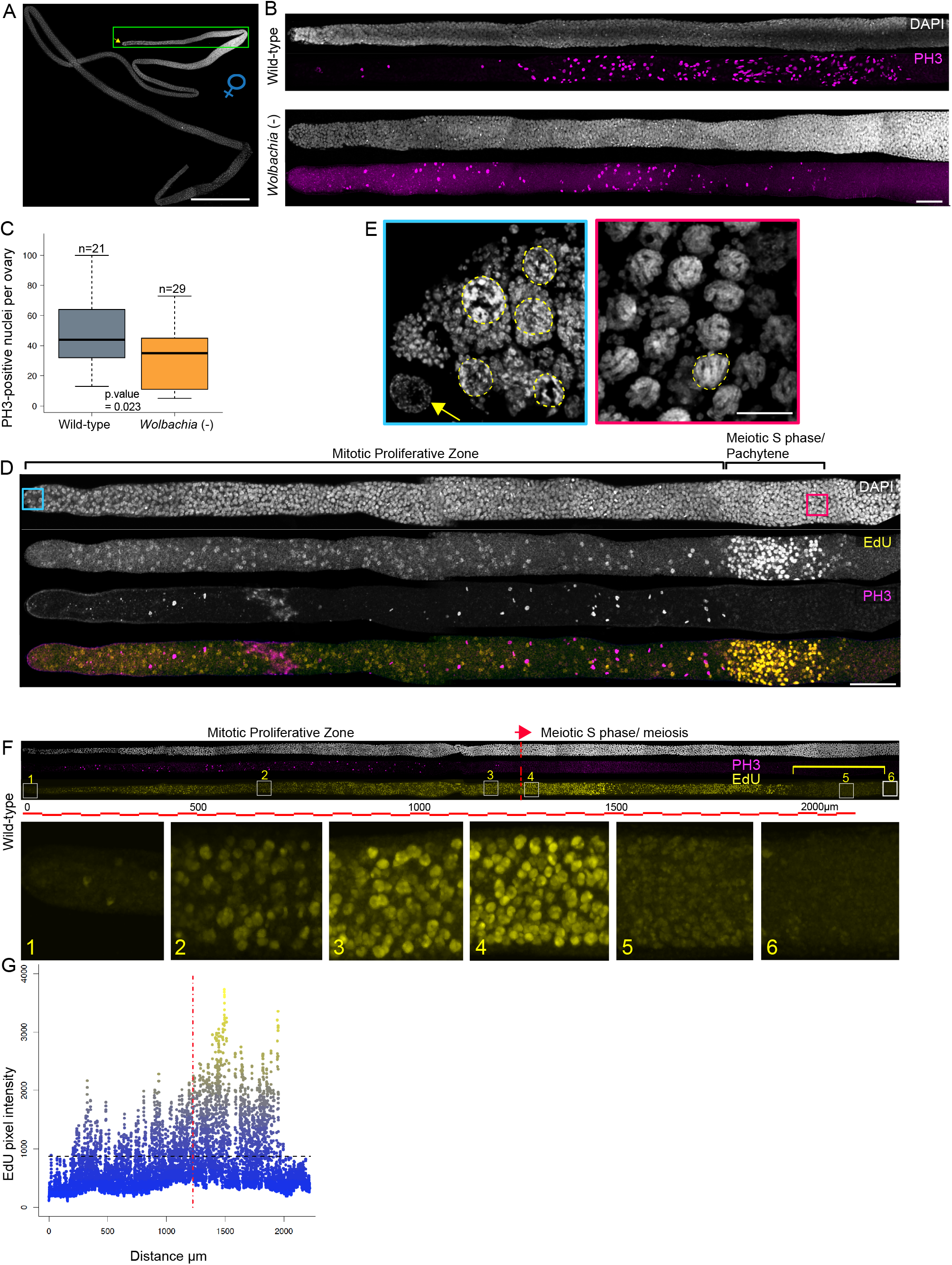
Effect of *Wolbachia* depletion on the germline proliferation and characterization of the proliferative zone in the ovary of *B. malayi*. Confocal images of (A) A dissected ovary stained with DAPI in its multiply folded natural conformation. The yellow arrow indicates the DTC. The narrow oviduct ends the ovary - bottom right-.(B) Full Z projection of confocal images of digitally linearized distal ovary parts from wild-type and *Wolbachia*-depleted females, stained to reveal DNA -DAPI in grey-, and mitotic nuclei - phosphorylated histone H3, "PH3" in magenta. Scale bar = 50 μm. (C) Quantification of the mitotic proliferation expressed as the number of phospho-histone 3 - positive nuclei –PH3 in 21 wild-type and 29 *Wolbachia* (-) ovaries respectively. Bold line: median; box: lower and upper quartiles; whiskers: smallest and largest non-outlier observations. (D) Z projection of confocal images of a digitally linearized distal ovary part corresponding to the green box in (A), from a female incubated in EdU for 8 hours and stained to reveal DNA -DAPI, grey-, EdU incorporation -yellow- and mitotic nuclei - PH3, magenta-. (E) Enlargements of DAPI-stained areas indicated in (D). In the blue box are shown nuclei -i.e. outlined in yellow-close to the DTC -yellow arrow-, and in the red box nuclei in the pachytene stage where synapsed chromosomes are visible. Foci revealed by DAPI in between nuclei are *Wolbachia*. (F) Confocal images of a dissected ovary after a 72hr-long EdU incubation and stained with the anti- PH3. Color codes as in (D). The position of the insets along the ovary are indicated by white boxes. (G) Quantification of the EdU fluorescence intensity along the ovary presented in (F). The black dotted line corresponds to the maximum intensity threshold for one complete round of replication, the red dotted line separates the PZ from the entry into meiosis as indicated in (F). Scale bars = 500 μm (A), 50 μm (B,D), and 5 μm (E).

### Germline nuclei in mitotic proliferative and meiotic differentiation zones are physically separated

Because of the similarities in gonad organization between nematodes belonging to the Secernentea, we established parallels between *B. malayi* and the experimental organism *C. elegans* whose germline development has been extensively studied (Hansen and Schedl, 2013). In both species the somatic gonad is organized as a tube capped by a distal tip cell - DTC- acting as a germline stem cell niche (Figure S1). In *C. elegans*, the DTC signals to germline nuclei, organized as a syncytium, to proliferate. These nuclei then exit the phase of mitotic proliferation and commit to the meiotic differentiation, initiated with a meiotic S phase. To characterize the *B. malayi* proliferative zone (PZ), worms were *in vitro* incubated with the thymidine analog EdU to reveal nuclei in S phase. Gonads were subsequently stained with an anti-PH3, and mounted with a DNA dye to identify the zone of meiotic entry (Figure 2D). The distal mitotic PZ corresponds to about one sixth of the entire ovary, and contains an average of 3000 nuclei (Figure 2A). Its proximal boundary was defined by the last PH3+ nucleus and the first EdU+ nuclei from the meiotic S phase cluster (Figure 2D). We found that in *B. malayi* females and males, nuclei in meiotic S phase are physically separated from nuclei in proliferation. This sharp transition to the meiotic S phase despite a high number of nuclei in the PZ suggests a tight spatiotemporal regulation of the differentiation mechanisms. These nuclei in S phase overlap with nuclei in the pachytene stage, in which synapsed chromosomes are clearly visible (Figures 2D and E; Figure S2).

### The germline undergoes transit amplifying divisions in the proliferative zone

To establish how the germline divides along the PZ, wild-type worms were exposed to EdU for 72 hours in *in vivo* conditions, in order to reveal altogether replicative and quiescent nuclei. Only proliferative nuclei in the PZ and those in meiotic S phase incorporate EdU. Therefore the number of EdU+ nuclei beyond the meiotic S phase area gives access to the kinetics of germ cells production. We localized the meiotic S phase area based on several criteria -its maximum EdU incorporation concomitant with nuclei displaying a chromatin typical of the pachytene stage, following the last PH3+ nuclei - (n>10, Figure 2F). The number of postmitotic EdU+ nuclei released from the meiotic S phase allowed us to estimate a yield of at least 700 germ cells produced per day per ovary. To understand whether they mainly arose from dispersed and actively dividing germline stem cells –GSCs- along the PZ, or from a distal pool of GSCs giving rise to daughters in transit amplification, we used the EdU fluorescence level to explore the germline nuclei cycling. To this end, we set up the imaging conditions using as a detection level threshold the fluorescence associated with the most proximal meiotic EdU+ cluster (Figure 2F, yellow bracket). This cluster reflects a single round of replication, undergone during the meiotic S phase only (Figure 2F inset #5). We found the distal area of the ovary to contain mostly EdU-negative cells, suggesting the presence of a quiescent pool of GSCs (Figure 2F inset #1). The fluorescence increase was then gradual along the 1mm-long PZ, from a basic level corresponding to a first round of replication and more (inset #2) to higher levels (insets #3 and #4). The quantification of the fluorescence intensity confirmed that the amount of incorporated EdU in the PZ goes beyond a single replicative cycle (Figure 2G). Yet the level of EdU incorporation appeared variable between nuclei close to the PZ exit (inset #3). Altogether these data suggest that i) the distal part is enriched in quiescent GSCs; ii) nuclei commit to differentiation in the proximal PZ after a variable number of transit amplification rounds.

### *Wolbachia* control the size of the proliferative pool in a cell-autonomous manner

We then explored the impact of *Wolbachia* depletion on the PZ, with the tools introduced above. Consistent with the reduced number of mitotic nuclei in absence of endosymbionts, we observed an average decrease of both the PZ length and its nuclear content by 33% compared to the wild-type (Figures 3A and 3B). We next wondered which of the soma or germline populations of *Wolbachia* is crucial to ensure a proper pool of proliferative nuclei. Since the antibiotic regimen wipes out *Wolbachia* from the somatic hypodermis of both female and male worms, but only from the female germline because the testis is devoid of endosymbionts, we reasoned as follows: If the proliferation of the male germline happens to be affected upon *Wolbachia* depletion, the somatic *Wolbachia* are very likely to participate to the germline proliferation. Conversely, if the male PZ does not appear affected, the role of somatic *Wolbachia* is probably insignificant in this regard, while the endosymbionts present in high titer in the female germline are very likely to be responsible for the enhanced proliferation. Comparison of germlines from wild-type versus *Wb*(*-*) males supports the latter. Neither the PZ length nor the number of nuclei in the testis PZ were affected (Figures 3C and S2C). To further confirm an overall absence of defects in spermatogenesis upon *Wolbachia* depletion, we followed the meiotic divisions and the sperm maturation in *Wb*(*-*) males and found no difference with the wild-type counterparts (Figure S2D). Incidentally, this cellular analysis shows that the tetracycline treatment itself does not directly cause germline defects, and altogether these data strongly suggest that *Wolbachia* support the female germline proliferation in a cell-autonomous manner.

**Figure 3.**
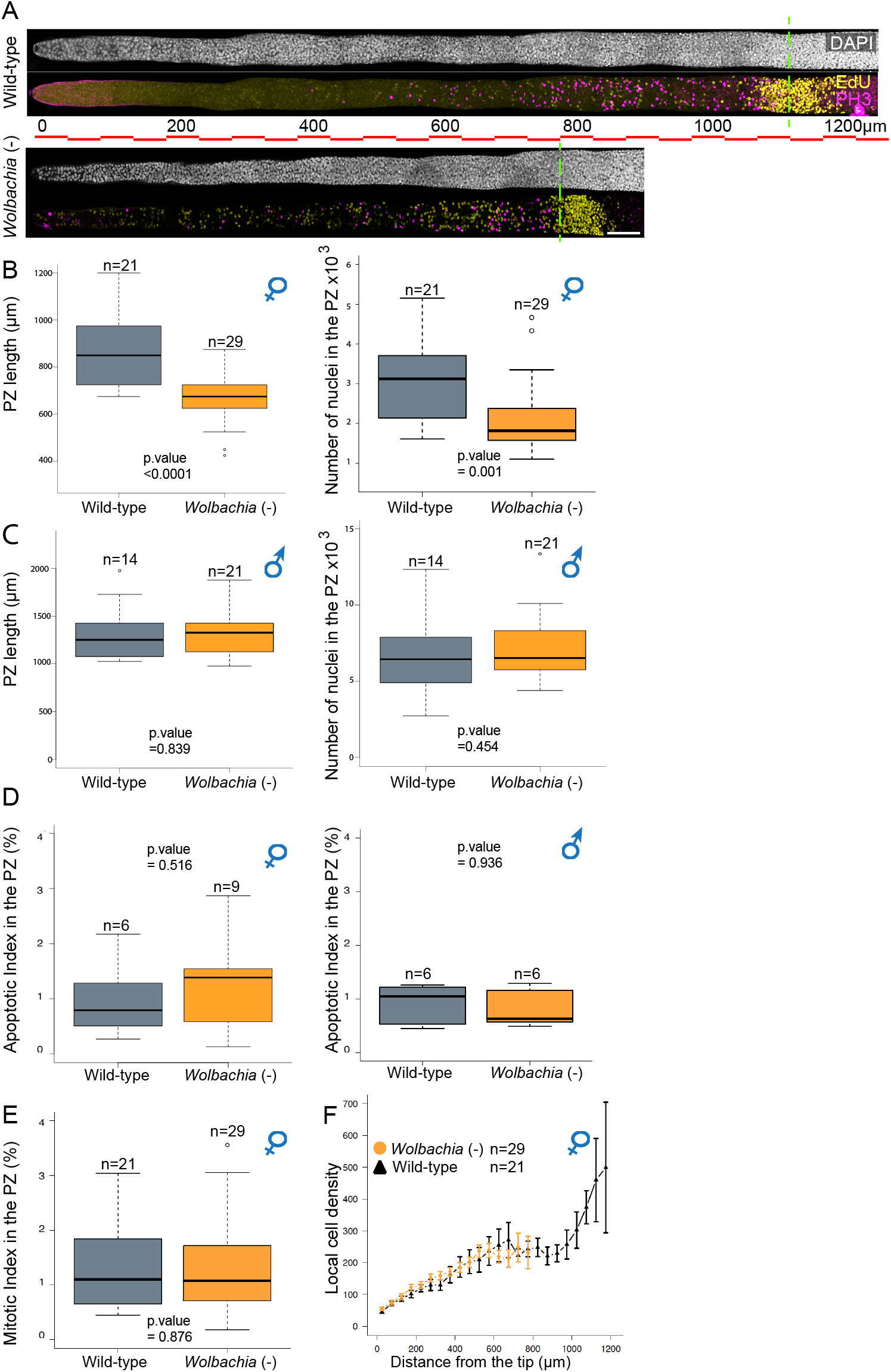
*Wolbachia* depletion affects the *B. malayi* female germline proliferative zone, in an autonomous manner. (A) Distal part of gonads dissected from wild-type and *Wolbachia*-depleted *B. malayi* females, stained with DAPI -grey-, and EdU -yellow- as well as PH3 -magenta-. The green dotted lines mark the limit of the PZs. Scale bar on images and scale bar unit on the ruler = 50 μm. (B) Length of the PZ and nuclear counts in wild-type and *Wolbachia-depleted* conditions in ovaries -top graphs- and (C) testes -bottom graphs-. (D) Apoptotic indexes in the PZ of ovaries -left- and testes -right- from wild-type and *Wolbachia-depleted* worms. (E) Mitotic index in the ovarian PZ of wild-type and *Wolbachia-depleted* females. For (B) to (E), bold line: median; box: lower and upper quartiles; whiskers: smallest and largest non-outlier observations; dots: outliers. (F) Density of nuclei by segments of 50μm along the ovaries of wild-type and *Wolbachia-depleted* worms.

### *Wolbachia* determine the transit amplification strength

Several mechanisms, not necessarily mutually exclusive, could account for a reduced number of mitotic nuclei in the female PZ, including an increase in distal apoptosis, a slower cell cycle, or less rounds of division associated with a precocious switch to meiotic differentiation. First, we scored the apoptotic nuclei in the PZs as the typically small condensed, PH3-negative pyknotic nuclei (Figures 3D and S3A). The comparison of apoptotic indexes of wild-type versus *Wb*(-) ovarian PZs showed no significant difference, and the same conclusion was reached for the testes (Figure 3D). Since the data were collected from worms sacrificed 8 weeks after the beginning of the antibiotic treatment of the rodent host, we envisaged that apoptosis might have occurred earlier and i.e. reduced the pool of GSCs. We therefore examined PZs at day 4 of host treatment, since apoptosis was previously reported in proximal ovaries and uteri at 4 days of *in vitro* treatment (Landmann et al., 2011). No significant apoptosis was detected (Figure S3B), and we concluded that the number of germline nuclei in the PZ is not reduced by an increase of cell death.

Second, a slower cell cycle in the PZ could in theory explain a smaller pool of PH3+ proliferative nuclei, shifting the balance toward differentiation. The mitotic indexes of wild-type and Wb(-) PZs appeared however similar, suggesting that proliferative cells are likely to cycle with the same speed (Figure 3E).

Last, the density of nuclei along the PZ was carefully examined. An estimate per segments of 50 μm revealed a constant average increase, similar in wild-type and *Wb*(-) PZs (Figure 3F). However the nuclear density in the last proximal segments of wild-type PZs showed a sharp increase in cell counts, suggesting a zone favorable to transit amplification. This zone is absent from Wb(-) PZs. Because the PZ lengths are naturally variable among same age females, we also scored the nuclear density as well as the distribution of mitotic events using heat map representations of each individual PZ (Figures S4A-S4B). These analyses confirmed the correlation between mitotic events and nuclear densities, and the increasing occurrence of mitotic events towards the proximal end of the wild-type PZs, clearly affected in *Wb*(-) PZs. Taken together, our data indicate that the absence of *Wolbachia* from the female germline does not result in distal apoptosis, nor in a change in the global cell cycle speed. Rather, the truncation of the transit amplification suggests that a role for the endosymbionts is to enhance the germline proliferation by increasing the mitotic cycling in the proximal part of the proliferative zone.

### The *B. malayi* female germline proliferation results from both Notch signaling pathway and *Wolbachia* inputs

Since *Wolbachia* modulate the level of germline proliferation, we wondered whether the symbionts operate through subversion of a host signaling pathway promoting the proliferation, or in parallel. In *C. elegans*, both specific developmental regulators as well as more ubiquitous cell cycle regulators have been identified to play key roles to sustain the proliferation (Crittenden et al., 1994; Fox et al., 2011; Lee et al., 2016). However most of the proliferation is under the control of a Notch pathway whose strength determines the PZ size (Austin and Kimble, 1987; Kimble, 2005). The transmembrane DSL ligand expressed by the DTC is delivered through long cell processes to the distal germline. Upon activation, the transmembrane Notch receptor is cleaved and translocated to the germline nuclei, promoting the expression of FBF RNA-binding proteins which prevent the meiotic differentiation (Lamont et al., 2004). We first explored the role of the DTC in the proliferation control by performing its laser ablation (Figure S5A). Only one DTC per female was ablated prior to comparative pair analyses. The count of PH3+ nuclei 24 hrs after DTC ablation showed no differences between ovarian pairs (Figures S5B-D) suggesting that in filarial nematodes the DTC is dispensable for the germline proliferation. This prompted us to assess the evolutionary conservation of the Notch pathway involvement in the germline proliferation control. To this end drug inhibition assays were carried out. *B. malayi* females were subjected to the potent gamma secretase inhibitor DBZ for 24 hours *in vitro*, that acts by preventing the cleavage of the Notch receptor (Fuwa et al., 2007). PH3+ nuclei counts revealed that although the loss of proliferation increased with higher doses of inhibitor (Figure 4A grey box plots), proliferation persisted specifically in the most proximal PZ part (Figure 4B wild-type). These cycling nuclei could either represent the last Notch-dependent proliferative nuclear pool, in which Notch downstream effectors are still active after the 24hr-long drug treatment, or a Notch-independent cycling. To test if this proximal cycling is promoted by *Wolbachia*, we exposed in parallel *Wb*(*-*) females to the same drug regimen. In that case, the germline proliferation was almost completely abrogated compared to the wild-type with a 100μΜ DBZ treatment (Figure 4A orange box plots, and Figure 4B), suggesting a *Wolbachia*-dependent proliferation control in the proximal PZ. To check if *Wolbachia* promote proliferation through a modulation of the Notch signaling pathway or independently, we identified - by reciprocal BLASTs with *C. elegans-* putative orthologs for key players of this pathway, and we measured the transcripts levels in wild-type versus *Wb*(*-*) ovaries by quantitative PCR (Figure 4C). We failed to detect any changes, supporting the idea that *Wolbachia* act independently of the Notch signaling pathway to promote proliferation. We then extended the qPCR analyses of ovaries to the orthologs of cell cycle regulators known to support the germline proliferation in *C. elegans* (i.e. *Bma cycline E, cdk2* and *cdc25*, see Figure S6 and Table S1). Again, no significant changes in expression levels of genes were detected. Altogether, this set of data indicates that i) a Notch ligand is likely to be expressed by other somatic cells than the DTC, ii) a Notch signaling pathway does control germline proliferation in *B. malayi*, iii) *Wolbachia* enhance the proliferation independently of the Notch pathway or other tested positive regulators of the germline division.

**Figure 4.**
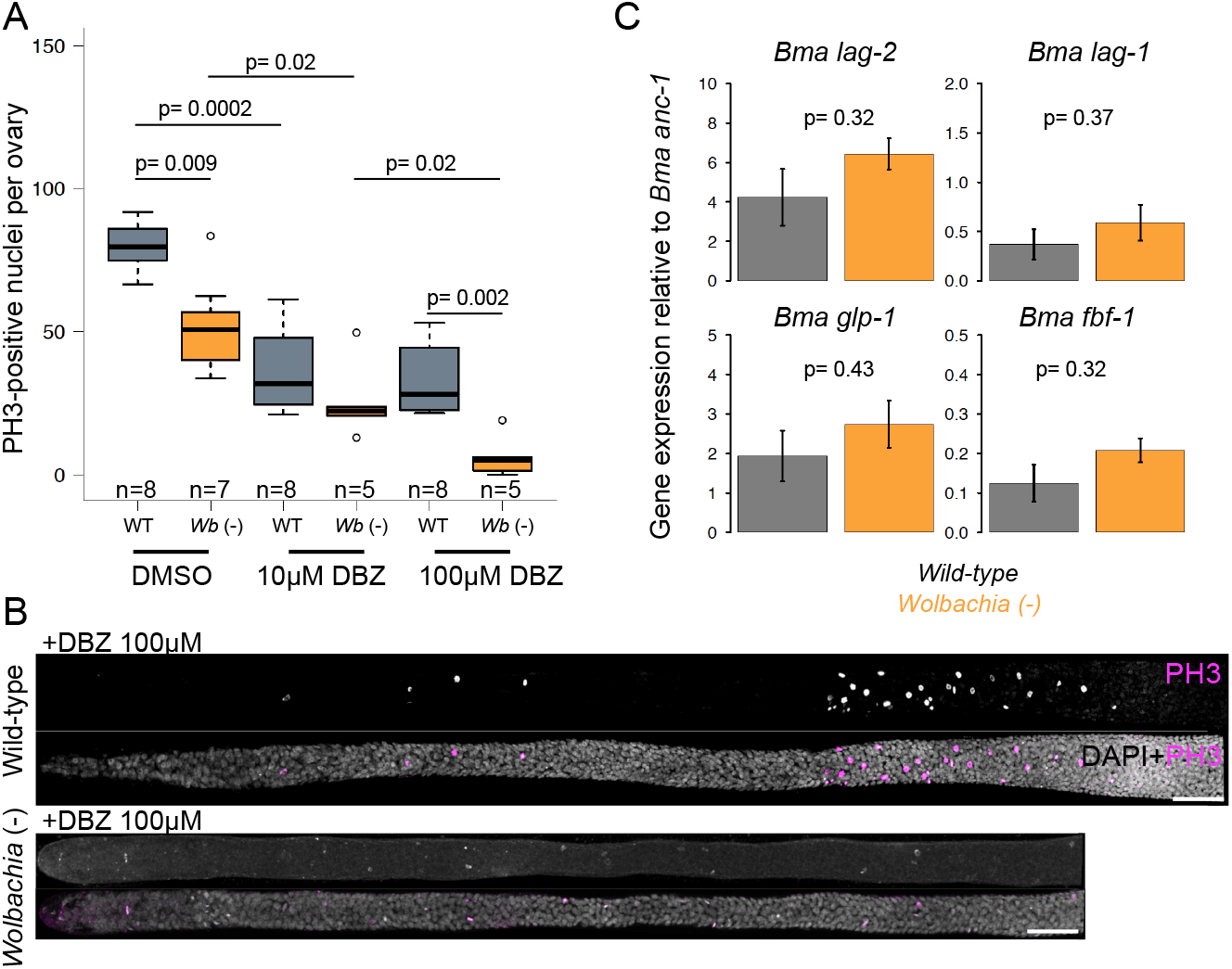
*Wolbachia* promote proliferation independently of the Notch signaling pathway. (A) Number of PH3+ nuclei counted in PZ from *B. malayi* females, wild-type or *Wb-*depleted, incubated for 24 hrs with either DMSO -control-, 10 or 100M DBZ. (B) Confocal images of distal ovaries from wild-type or *Wb-*depleted females incubated for 24 hrs in 100 μm DBZ, stained with DAPI -grey- and PH3 -magenta- (C) qPCR experiments on *B. malayi* genes whose orthologs are involved in the *C. elegans* Notch signaling pathway (*lag* ligands; *glp-1* receptor, *fbf-1* a Notch pathway downstream target).

### *Wolbachia* depletion does not induce a critical shortage of nucleotides

Since we did not identify any proliferation key players to be affected by the loss of *Wolbachia*, we looked at the nucleotide levels. Cell proliferation demands an important nucleotide supply, and the available pyrimidine pool of nucleotides was recently shown to be critical to sustain the germline proliferation in *C. elegans* (Chi et al., 2016). In addition, a genomic analysis of *Wbm Wolbachia* indicates that these intracellular bacteria have retained the metabolic capabilities to *de novo* synthesize nucleotides (Foster et al., 2005). To test the hypothesis of this metabolic contribution as one of the possible bases of the mutualism with the worm, nucleotides levels were measured in female worms. Because of the large amount of bacteria in somatic and germinal tissues, we hypothesized that the symbionts depletion could *de facto* induce a corresponding decrease in global nucleotide pools. We therefore added a filarial nematode species naturally devoid of *Wolbachia, Acanthocheilonema viteae*, as a control (Figure 5). While ATP and ADP levels were above the upper threshold in our calibration conditions to be reported, a significant decrease of all other nucleotides was rarely measured between wild-type and *Wolbachia-depleted B. malayi* females. Whenever observed (i.e. for GMP and GDP), the pools were similar between *Wb*(*-*) *B. malayi* and *A. viteae*, suggesting that these concentrations are enough to sustain fertility and survival. Moreover, the variations never exceeded a two-fold range, while pathological states associated with nucleotide metabolism defects present much higher disturbances in the nucleotides levels (Bester et al., 2011; Mathews, 2015). Altogether, these data suggest that the mutualism between *Wolbachia* and the filarial host is unlikely to involve a nucleotide supply of bacterial origin.

**Figure 5.**
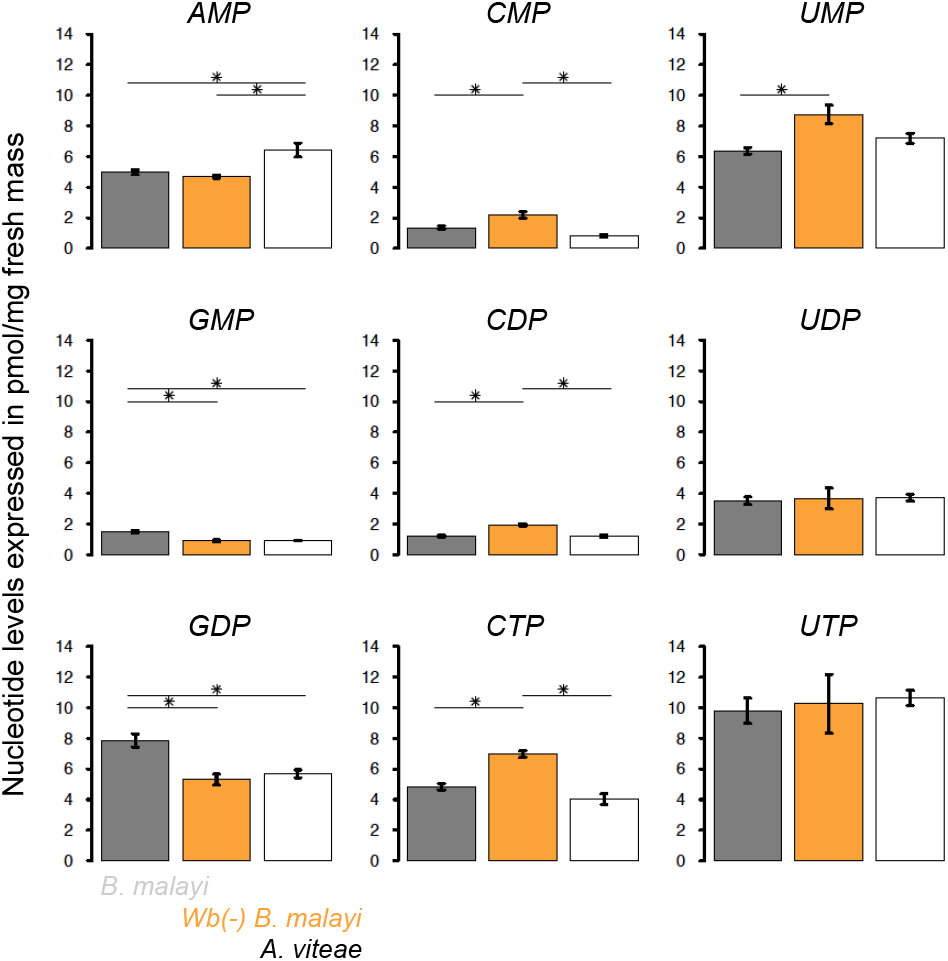
Influence of *Wolbachia* on the nucleotide levels. Amounts of various nucleosides mono-, di-, and triphosphates, expressed in pmol/mg of fresh female worms, in wild-type *B. malayi* -grey-; *Wolbachia-depleted B. malayi* -orange-; or *A. viteae* -white-species.

### *Wolbachia* control germline stem cells behavior and organization

We wondered if *Wolbachia* influence would also extend to the GSCs. Although there are no specific markers for this cell type in nematodes, we took advantage of the identification of a quiescent pool of distal cells, *de facto* GSCs, spanning over the first 150 μm (Figures 1 and S4). To check a possible effect of *Wolbachia* on the GSC quiescence, and because of the rare occurrence of mitotic events in this zone, we incubated wild-type and *Wb*(*-*) females with colchicine, in order block and accumulate mitotic events (Figure 6A). We indirectly characterized the quiescence by scoring the PH3-positive nuclei over the most distal 150μm, in three adjacent 50 μm-long segments (Figures 6A and 6B). We found in wild-type ovaries a gradient of division events, indicating the presence of more quiescent nuclei towards the distal tip. A laser ablation experiment of the DTC followed by a colchicine incubation ruled out a role of this cell in the quiescence control (Figure S7A). The observation of *Wb*(*-*) ovaries revealed that the GSCs divide on average 4.77 times more along the first 50 μm in absence of *Wolbachia* than in control ovaries, twice more in the second segment, and 1.41 more in the last segment. Additional *in vivo* EdU incorporation experiments revealed similarly an average fold change of GSCs in S phase of respectively 4.03; 2.95 and 1.32 in the same three segments in *Wb*(*-*) ovaries compared to wild-type ovaries (Figures S7B and S7C). Altogether these data indicate that the germline stem cells quiescence is controlled by *Wolbachia* in *B. malayi*.

**Figure 6.**
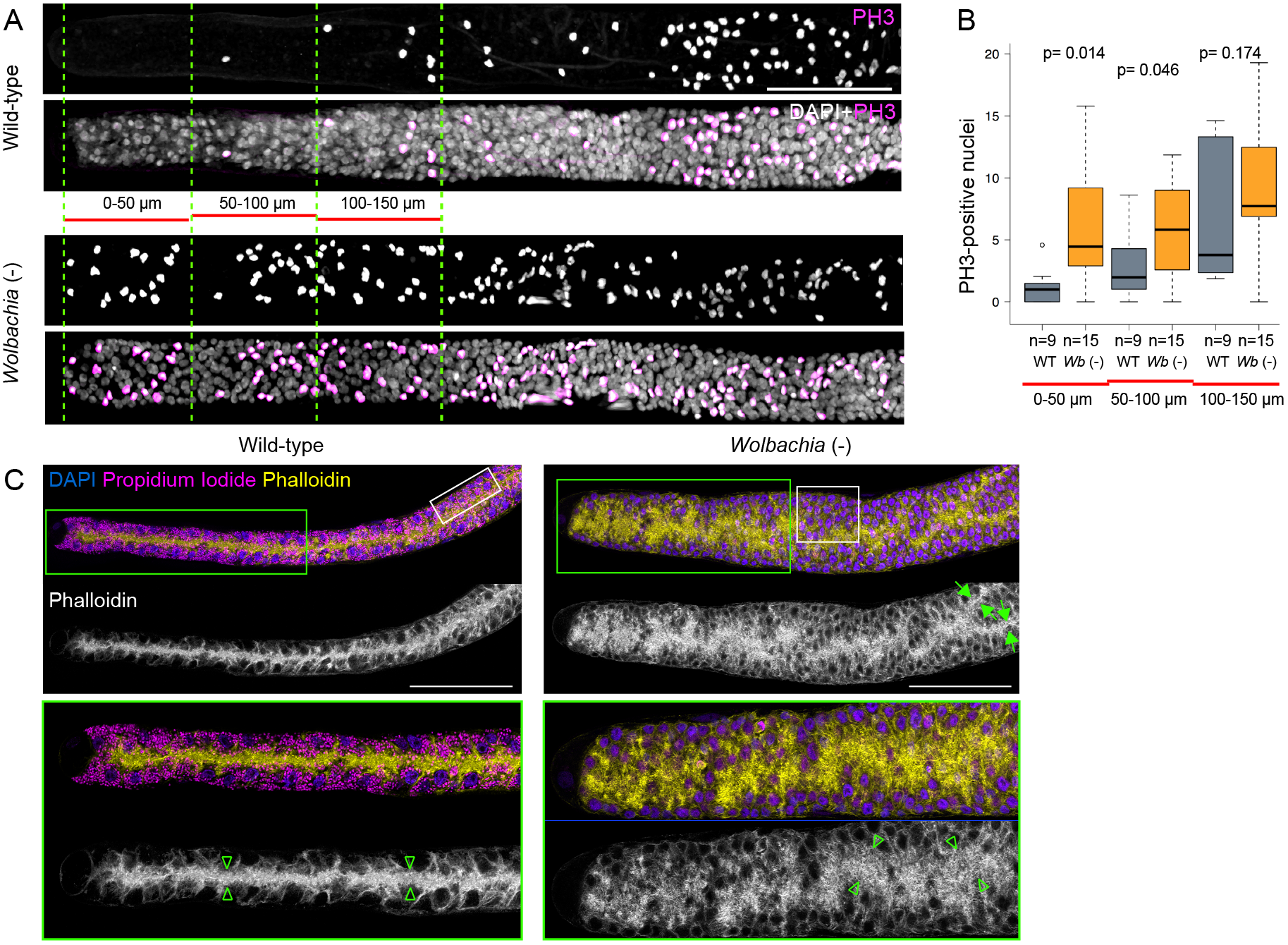
The loss of *Wolbachia* modifies germline stem cells features. (A) Confocal acquisitions of digitally linearized distal ovaries from wild-type and *Wolbachia-depleted B. malayi* females incubated *in vitro* for 4 hours in presence of colchicine 1mM, and stained with DAPI -grey- and an anti-PH3 -magenta-. The 50 μm long segments are marked by a green dotted line. Scale bars = 50 μm. (B) Number of PH3-positive nuclei counted in the most distal 0 to 50 μm and 0 to 150 μm-long segments of ovaries from wild-type and *Wolbachia-depleted* females. (C) Distal parts of ovaries double stained for DNA (with DAPI - blue-, staining mainly the host nuclei, and propidium iodide -magenta-revealing the *Wolbachia-*) and phalloidin -yellow or grey-decorating the actin cytoskeleton. Lower panels are enlargements of areas marked in green. White boxes highlight the nuclear organization around the rachis. Green arrows point to a branched rachis. Green arrowheads encompass the rachis.

Last, we explored if other GSC behavioral traits could be affected by the loss of the endosymbionts. We took a closer look at the rachis and the distal nuclear distribution. The germline syncytium is organized as a single layer of cortical nuclei forming an actin-rich central canal in wild-type PZs (Figure 6C). The absence of *Wolbachia* affected the rachis structure that appeared very often disorganized, larger, and branched. This was correlated with a disorganization of the distal nuclear layer (i.e. Figure 6C, white inset in *Wb*(*-*) distal part), indicating that the GSCs behavior was perturbed. We concluded that the *Wolbachia* endosymbionts are necessary to achieve a correct germline stem cell developmental program in the filarial host.

## DISCUSSION

In this study, we demonstrate that a germline development goes awry without the presence of intracellular bacteria. In filarial nematode species living in mutualism with *Wolbachia*, both the long-term survival and the fertility of the host depend on the endosymbionts. Although they can live for months without *Wolbachia*, the infertility occurs following the endosymbionts depletion (Hoerauf et al., 1999). While it was previously established that the *Wolbachia* depletion eventually triggers apoptosis during embryogenesis and sometimes during late oogenesis in proximal ovaries (Landmann et al., 2011), nothing was known about the influence of these bacteria during early oogenesis. We circumvented technical obstacles to characterize the early oogenesis of these unwieldy nematodes. This allowed us to define with a fine scale spatiotemporal resolution the contribution of *Wolbachia* to discrete events during the germline proliferation in *B. malayi* (summarized in Figure 7). Specifically, we found the most distal GSCs to be in a quiescent state that depends on *Wolbachia*. This asymmetric enrichment of quiescent stem cells in the PZ suggests that the increasing number of mitotic events along the PZ reflects a transit amplification resulting mainly from the division of a distal pool of active GSCs. While the proliferation partially depends upon a Notch signaling pathway, our results show that the *Wolbachia* present in the germline enhance the mitotic events in a cell autonomous manner. In addition, the distal-most defects induced by the loss of *Wolbachia* suggest an active participation of the endosymbionts in the proper maintenance of the female germline stem cell fate.

**Figure 7.**
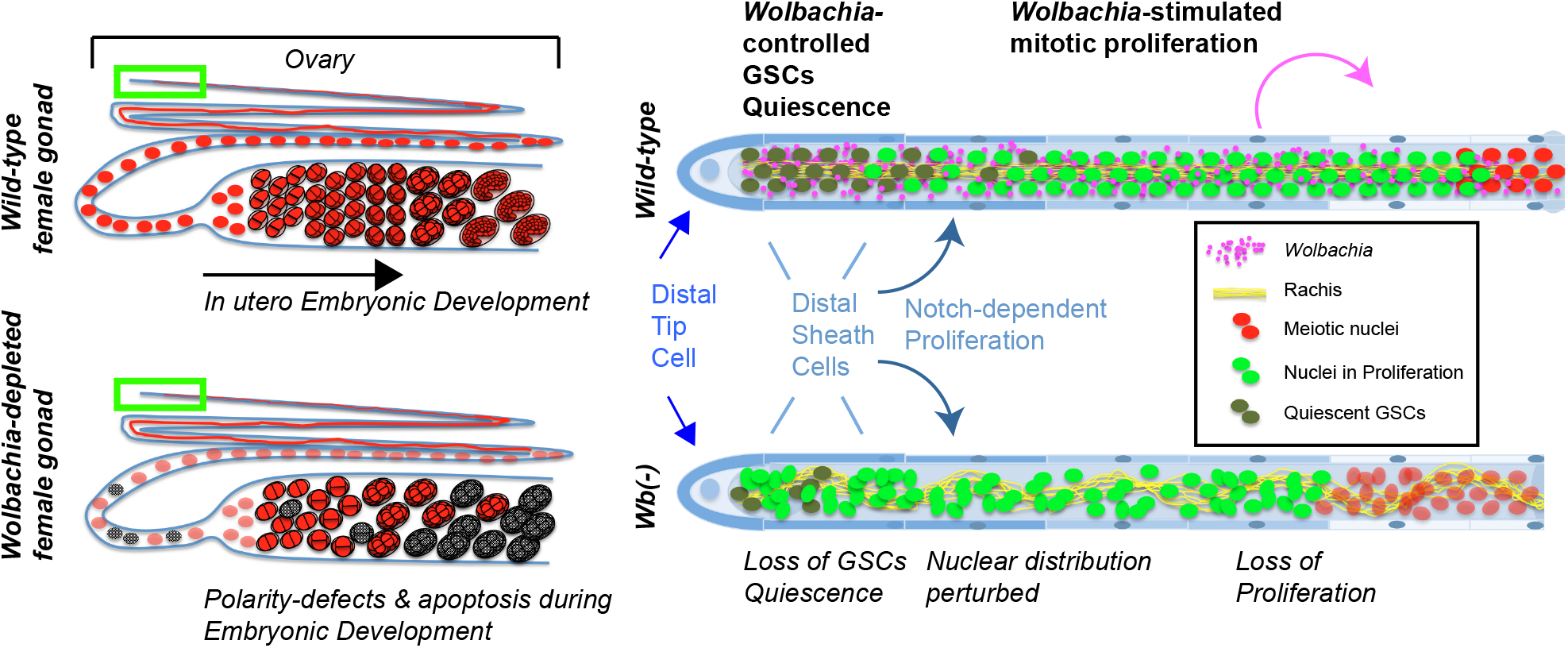
Contribution of *Wolbachia* to the *B. malayi* oogenesis and phenotypes resulting from the endosymbionts depletion. Cartoons on the left are schematic representations of full length ovaries connected to distal uteri. They depict the early embryonic development, altered in absence of *Wolbachia* (i.e. apoptosis in grey). The green rectangles highlight the distal ovaries represented on the right.

Understanding how *Wolbachia* influence the germline implies to know to some extent their host's peculiar biology. Filarial nematodes and *C. elegans* differ by their lifestyles and are phylogenetically distant enough (parasitic from clade III, and free-living from clade V respectively (Blaxter et al., 1998)) to consider the latter one only as a starting point. This study reveals unique features of *B. malayi* oogenesis related to its parasitic lifestyle. We estimate a female to produce about 1400 eggs per day, resulting in as many microfilariae released in the blood of the vertebrate host on a daily basis. These long-lived worms can actively reproduce for 5 to 8 years (Taylor et al., 2010). A lower estimate of 1.3 million eggs per ovary is therefore produced during a lifetime. To face such a demand, *B. malayi* possesses a gigantic proliferative zone made of about 3,000 nuclei. If these nuclei were all equivalent in proliferative properties, each would divide more than 400 times. While the very short-lived *C. elegans* counts only 200 to 250 actively cycling nuclei in the PZ (Crittenden et al., 2006), the presence in *B. malayi* of hundreds of GSCs in quiescence associated with a transit amplification phase is likely to reflect an alternative strategy to protect the germline. It is indeed crucial for stem cells to prevent accumulation of DNA replication-dependent mutations. To this end, only a fraction of GSCs transiently divide to give rise to transit amplification cells, likely to be gradually replaced by fresh stem cells from the quiescent distal pool. Which cells control the proliferation and the quiescence is not clear yet. In *C. elegans*, the DTC promotes the gonad migration and morphogenesis during larval development (Wong and Schwarzbauer, 2012), in addition to serve as a niche producing the Notch ligand controlling the proliferation in the adult germline (Austin and Kimble, 1987; Kimble and White, 1981; Lee et al., 2016). Our laser ablation experiments show that in *B. malayi* the germline proliferation does not depend on the DTC. Yet its reliance on a Notch signaling revealed by a gamma-secretase inhibition assay indicates that other somatic cells must provide the ligand. While the *C. elegans* DTC has been proposed to deliver the Notch ligand through cell processes forming a plexus running in between the distal germline (Byrd et al., 2014), the scale of oogenesis in filarial nematodes may rely on broader somatic sources of signal, i.e. the distal sheath cells. In fact, *C. elegans* sheath cells show different properties along the gonad and the most distal ones participate to the larval germline proliferation (Killian and Hubbard, 2005; McCarter et al., 1997). Moreover It was shown that the *Wolbachia* tropism for the *Brugia* germline is limited to the most distal sheath cells, through which the symbionts enter the gonad, suggesting different somatic cell properties along the gonad (Landmann et al., 2012). It is likely that in adult *B. malayi*, distal sheath cells support proliferation, while the main function of the DTC would remain to lead the gonad migration during development. We established that the DTC does not control the GSC quiescence. It also seems unlikely that a Notch signaling pathway would regulate quiescence since in *C. elegans* facultative stem cell quiescence induced by food restriction is independent of Notch (Seidel and Kimble, 2015).

Being vertically transmitted through the female germline, the *Wolbachia* have become master manipulators of their hosts' reproduction machineries to ensure their transmission to the next generation and invade a host population (Zug and Hammerstein, 2014). Facultative *Wolbachia* have been shown to stimulate the egg production in a *Drosophila* species by enhancing the GSCs division together with an inhibition of apoptosis in the germline (Fast et al., 2011). In that case, their presence in the GSC niche was proposed to enhance GSCs division through a non cell-autonomous mechanism. We describe here a different mechanism of germline division enhancement. The *Wolbachia* living in mutualism with *B. malayi* do not target the DTC, and are absent or poorly present in the surrounding distal somatic sheath cells. Rather, they seem to act in a cell-autonomous manner in the female nematode germline. They do not influence the proliferation by a modulation of apoptotis levels in the PZ. Upon their depletion, a third of the cycling nuclei is lost, reflected accordingly in the number of nuclei in the PZ. The mitotic index is therefore not influenced by the presence of the symbionts, suggesting that *Wolbachia* are unlikely to modify the cell cycle.

Two scenarios not necessarily mutually exclusive, and developed hereafter, could explain how *Wolbachia* support the germline proliferation. First, *Wolbachia* could participate to the GSC fate maintenance. Second, the symbionts could stabilize the mitotic proliferation stage and delay the commitment into meiosis.

In absence of *Wolbachia*, the GSC quiescence is heavily perturbed, demonstrating a clear role of *Wolbachia* in its maintenance. Beyond the observed quiescence alteration, *Wolbachia* depletion could induce more global defects in GSC fate maintenance, that would explain not only the ectopic divisions occurring in the most distal part of the ovary, but also the disturbed spatial organization of these nuclei correlated with a misshapen rachis. A reduction of GSC self-renewal upon division would lead to a reduction of the proliferative pool.

We also show that upon a drug-induced Notch signaling inhibition for 24 hours, a fraction of nuclei still proliferates in the proximal PZ. This remaining proliferation is lost when the *Wolbachia* are removed. Although our candidate approach based on orthologs of key controllers of the *C. elegans* germline proliferation failed to identify a potential host signaling mechanism by which *Wolbachia* could enhance this proximal proliferation, it however suggested that the symbionts act in parallel of the Notch signaling pathway.

The still mysterious interdependency between *B. malayi* and its symbiotic *Wolbachia* strain *Wbm* has been hypothesized to rely on the metabolic capabilities of each partner (Fenn and Blaxter, 2007; Foster et al., 2005). *Wbm* have retained complete pathways to synthesize purines and pyrimidines, and could potentially supplement the host pool. We tested their relevance in the ovary, since i) the germline is the only proliferative tissue in the adult female, in addition to the embryogenesis, both demanding in nucleotides, and ii) while only a handful of *Wolbachia* is found in developing embryos, limited to few specific blastomeres (Landmann et al., 2010), their titer is much greater in the ovarian PZ, and therefore more likely to exert an influence on the germline proliferation, shown to depend on the available pool of nucleotides in *C. elegans* (Chi et al., 2016). We did not find supporting evidence of such a dependency since nucleotide pools are not reduced in *Wb-*depleted females beyond their normal concentrations in *A. viteae*. The idea that synthesis pathways, missing or partially missing in the host would be complemented by *Wolbachia* has also been seriously undermined by the publication of the *Loa loa* genomics, a *Wolbachia-free* filarial nematode species (Desjardins et al., 2013). Despite the lack of the symbionts, *Loa loa* does not present additional metabolic pathways compared to *B. malayi*, suggesting more subtle mechanisms underlying the symbiotic association.

In *C. elegans*, the self-renewal of the GSCs together with the germline mitotic proliferation are maintained through Notch signaling-dependent mechanisms that essentially operate through post-transcriptional repression of meiotic differentiation regulators thanks to the RNA binding proteins FBFs (Crittenden et al., 2002; Lamont et al., 2004). All these regulators are conserved broadly and are very likely to act in the control of germline proliferation in filarial nematodes. Our data reveal that the *Wolbachia* depletion results in a global decrease of germline proliferation with a loss of GSC quiescence. The two phenotypes triggered by the depletion of *Wolbachia* (i.e. the global decrease of germline proliferation and the loss of GSC quiescence) may share the same origin. Defects in GSC fate maintenance would force the distal nuclei to exit quiescence and eventually would lead to a loss of self-renewal, reducing the pool of proliferative nuclei.

The addition of *Wmel Wolbachia* strain to their natural host *D. melanogaster* has been shown to rescue germline defects in *Sex-lethal* mutants by restoring GSCs maintenance through the effector TomO, acting at the *nanos* mRNA level to increase its translation and support the GSC fate (Ote et al., 2016; Starr and Cline, 2002). Although TomO is not conserved in *Wbm*, this strain also harbors a functional type IV secretion system releasing bacterial effectors in the host (Li and Carlow, 2012). We can speculate that *Wolbachia* may interfere at the translational level to participate to the repression of the meiotic differentiation. In addition, because all embryos before morphogenesis, and sometimes cellularized oocytes in the most proximal ovary eventually enter apoptosis, it is possible that *Wolbachia* became essential to ensure a correct oogenesis developmental program, leading otherwise to cell death. For instance, polarity defects observed in *Wb*(*-*) early embryos could results from earlier defects during oogenesis (Landmann et al., 2014). Intracellular pathogens have been shown to modulate host gene expression to their advantage through epigenetic modulations (Bierne et al., 2012). Whether this developmental symbiosis, leading otherwise in the absence of *Wolbachia* to defects from the GSCs to the early *B. malayi* embryo is under a symbiont-controlled epigenetic program will be the focus of future studies.

## AUTHOR CONTRIBUTION

V.F., M.M.P.J. and F.L. conceived experiments, performed formal analyses and data presentation. V.F., M.M.P.J. and N.F. performed experiments. F.V. and L.F. wrote the manuscript.

## ACKNOWLEDGMENTS

MetaToul (Metabolomics & Fluxomics Facitilies, Toulouse, France, www.metatoul.fr, MetaboHUB-ANR-11-INBS-0010) and its staff members are gratefully acknowledged for carrying out metabolome analyses. We are grateful to NIH/NIAID Filariasis Research Reagent Resource Center (www.filariasiscenter.org) for providing *B. malayi* and *A. viteae* specimens. We thank V. Georget and S. Lachambre from Montpellier RIO Imaging for microscopy facilities and their technical support for image analyses. We thank Pascale Cossart for critical advice. We are also grateful to Coralie Martin for advice on filarial nematode rearing techniques. This work was supported by the « Fondation pour la Recherche Médicale » (ARF20150934088) and the ATIP-Avenir program.

## STAR METHODS

### Contact for Reagent and Resource Sharing

Further information and requests for resources and reagents should be directed to and will be fulfilled by the Lead Contact, Frédéric Landmann

### Experimental Model and Subject Details

#### Ethics statement

All experiments involving animals were approved by the ethical review committee of the Ministère de l’Education Nationale, de l'Enseignement Supérieur et de la Recherche (authorization #03622.01). Housing, breeding and animal care were carried out in strict accordance with the EU Directive 2010/63/UE.

#### Parasite material

Living *B. malayi* worms, harvested from infected jirds (*Meriones unguiculatus*), were supplied by the NIAID/NIH Filariasis Research Reagent Resource Center (FR3, Athens, USA) via the Biodefense and Emerging Infections Research Resources Repository (BEI Resources, Manassas, USA) or produced at the Institut de Recherche pour le Développement (IRD, Montpellier, France). Jirds were infected by injection in the peritoneal cavity of 100-200 infective larvae (L3s) freshly collected from *Aedes aegypti* Strain Black Eye Liverpool, following the FR3 protocols (http://www.filariasiscenter.org/protocols/Protocols). Living *Acanthocheilonema viteae* worms, harvested from infected Golden Syrian LVG Hamsters (*Mesocricetus auratus*), were supplied by the NIAID/NIH Filariasis Research Reagent Resource Center (FR3, Oshkosh, USA). To obtain *Wolbachia-depleted B. malayi*, jirds received tetracycline at 2.5 mg/mL in drinking water 90 days post-infection (dpi) during a period of 6 weeks, or tetracycline at 50 mg/kg/day during two weeks *per os* using a solution at 10 mg/mL. Worms were collected from the peritoneal cavity two weeks after the end of the antibiotic treatment, and the *Wolbachia* clearance in the soma and germline confirmed by fluorescent microscopy. Wild-type counterparts were obtained from jirds maintained for the same duration without the tetracycline regimen. For *in vitro* live assays, living worms were placed in culture medium (80% RPMI-1640, 10% decomplemented FBS, 10% MEM, 1% glucose, 25 mM HEPES buffer, pH 7.4), in incubation at 37°C and 5% CO_2_. Typically, 0.5 or 1 mL/worm/day of culture medium were used for adult male and *female B. malayi* respectively, in 6 or 24-well culture plate (Greiner Bio-One), and the culture medium was changed every 48 hours. At the end of experiments, worms were flash frozen in liquid nitrogen and kept at -80°C until dissection.

### Method Details

#### Tissues collection

Frozen worms were thawed at room temperature and fixed in a 3,2% paraformaldehyde (Electron Microscopy Sciences, #15714) PBS solution during either 10 or 15 minutes on a rocker for males and females respectively, then washed 3 times in PBST (1x PBS, Tween-20 0,2%). Dissections were performed in PBST under binocular microscope (SMZ1270; Nikon) with dissection tweezers. To collect ovaries, a first incision was performed in the posterior part, close to the distal uteri, by gently tearing apart the body walls, without breaking the gonads and intestine. A second cut at the very end of the tail prevents internal pressure when pulling on the posterior body fragment. Both ovaries were then carefully pulled out of the body cavity. The same procedure was applied to collect the testis except that the first incision was performed at the half of the body and the second at the tip of the head since the distal part of the testis lies at the level of the pharynx.

#### Stainings

For immunostaining, dissected gonads were collected in 0.5 mL eppendorf tubes with PBST, and permeabilized with the following protocol empirically established: after a 45 minute incubation with a 3% hydrogen peroxide solution, gonads were washed three times in PBST, treated for 30 minutes with a (1:1) mix of heptane and NP40 2% in PBS, and then washed 3 times in PBST. All these steps were performed on a rocker at room temperature. Gonads were incubated overnight at 4°C with a primary antibody, washed 3 times in PBST and incubated overnight at 4°C or alternatively for 6h at room temperature with a secondary antibody, washed 3 times in PBST. Mitotic nuclei were revealed with an anti-phospho-histone 3 pSer 10 (PH3), rabbit monoclonal antibody (Invitrogen, # 701258, 1:250). Because tri-methylation on lysine 27 of histone 3 is enriched on X-chromosome in *C. elegans* during spermatogenesis (Schaner and Kelly, 2006), an anti-H3K27me3 rabbit polyclonal (Epigentek A-4039, 1:250), was used to monitored the correct chromosome segregation in male *B. malayi*. We used a goat anti-rabbit IgG secondary Cy3 conjugated antibody, (Invitrogen, #A10520, 1:250). Actin staining was performed using Phalloidin Rhodamine 110 conjugate (Biotium, 1:100), added with together with the secondary antibody. As previously described (Landmann et al., 2010, 2014), *Wolbachia* were specifically stained with a short incubation in propidium iodide (Invitrogen, #P3566, 10 μg/mL in PBS) after treatment with RNAse A (Sigma-Aldrich, #R6513, 10 mg/mL in PBS) overnight at 4°C. Samples were mounted in Fluoroshield Mounting Medium with DAPI (Abcam, #Ab104139).

#### Edu assays

For *in vitro* analyses, living adult worms were placed in culture medium preheated at 37°C supplemented with 200 μM 5-ethynyl-2´-deoxyuridine (EdU, Invitrogen, #A10044), using a stock solution at 10 mM in DMSO. After *ad hoc* incubation times, worms were flash frozen. EdU was revealed after fixation, dissection and permeabilization steps using the Click-It EdU Alexa Fluor 488 Imaging Kit (Invitrogen, #C10337) following the manufacturer’s instructions, except that reactions were performed in 0.5 ml tubes.

For *in vivo* assays, infected jirds received one intra-peritoneal injection of EdU per day during three consecutive days. EdU was given at a dose of 50 mg/kg body weight in a solution of 10mg/ml PBS (Chehrehasa et al., 2009) and the first injection was performed after 90 days post infection. Worms were collected from the peritoneal cavity 24 hrs after the last injection and immediately flash frozen. EdU Clik-It reactions were performed as described above. When EdU analyses were combined with immunostainings, the EdU Click-It steps were performed before the incubation with the primary and secondary antibodies.

#### Colchicine assays

Living adult worms were placed in preheated culture medium at 37°C supplemented with 1mM colchicine (Sigma-Aldrich, #C9754) using a stock solution at 100 mM in ethanol. Worms were incubated 4 hrs at 37°C and were then flash frozen. No effect of ethanol on the mitotic proliferation was observed in control experiments (data not shown).

#### Gamma-secretase inhibitor

A 10 mM stock of Dibenzazepine (DBZ, Stemcell Technologies, #73092) in DMSO was diluted in preheated culture medium at 37°C. Based on previous studies using DBZ (Ichida et al., 2014) or an other gamma-secretase inhibitor (Geling et al., 2002), DBZ has been tested at 10 and 100 μM. Control worms were mock-treated with medium containing the same concentration of DMSO carrier only.

#### Laser ablation

Living adult worms were individually immobilized in preheated culture medium at 37°C supplemented with 1 mM levamisol (Sigma-Aldrich, #31742) during few seconds and mounted on an agar pad (0.6% in PBS) under a coverslip. Before imaging, the slide was maintained on ice to maintain the anaesthesia. The laser microsurgery was performed using an Ultra II Coherent multiphoton laser at 800 nm full power coupled to a Zeiss LSM780 confocal microscope. The sample was imaged in transmission mode using the 561nm laser and the 63X 1.4NA oil immersion objective. The ablation to kill the cell was obtained with 1 to 10 iterations in a region of 10*10 μm. The distal tip cell (DTC) was ablated from one ovary per worm and the remaining ovary was kept intact as an internal negative control. When the targeted cell appeared to be destroyed, the worm was returned to *in vitro* culture maintained during 24 hrs before analysis. During all the subsequent steps, ovaries from the same female were processed together, and the pair information was taken in account in statistical analyses.

#### Microscopy and Image analyses

Confocal microscope images were captured with an inverted laser scanning confocal microscope (SP5-SMD; Leica Microsystems) using a 63X/1.4 HCX PL APO CS oil objective and a resonant scanner (8000Hz). Gonads were fully imaged with a z-stack interval of 0.5 μm and identical imaging conditions. The digital images were processed and analyzed using a custom macro with the ImageJ 1.48v software (http://imagej.nih.gov/ij/) to semi-automatically quantify the number and the distribution of cells *i*) in the proliferative zone, *ii*) in mitosis and *iii*) in S-phase. Briefly, the gonad was first digitally linearized with a straightening function and then analyzed per 50μm-wide sections starting from the distal tip cell. For each section, we estimated *i*) the total number of nuclei, by multiplying the sum of DAPI area from one median z-stack by the number of nuclei rows in the z-axis; *ii*) the number of mitotic and S-phase cells by summing the PH3 and EdU areas respectively, in the z-projection of all planes. To reduce the background noise, a size particle selection filter was manually applied before area summation to remove particles with an area three-time smaller or larger than the mean nucleus area. The accuracy of this procedure was compared to manual counts on few gonads and all data obtained with this procedure were manually curated.

#### Expression of candidate genes

Ovaries used for RNA extractions were dissected from living adult females. Dissections were performed under a binocular microscope as described above, in a RNAse-free environment by placing the worms in sterile PBS 1X and cleaning all the materials with RNAseZap (Invitrogen, #AM9780). Five pairs of ovaries were pooled per biological replicate in a frozen tube maintained on dry ice. Tubes were flash frozen and stored at -80°C until RNA extraction. Extractions were performed using the Quick RNA MicroPrep kit (Zymo-Research, #R1050) and residual contaminant DNA was removed with Turbo DNAse (Invitrogen, #AM1907), followed by purification using the RNA Clean & Concentrator 5 (Zymo-Research, #R1016). The RNA yields were determined fluorometrically using Qubit 2.0 (Life Technologies). The cDNA was synthesized using the SuperScript^®^ VILO cDNA Synthesis Kit (Invitrogen, #11754050), according to manufacturer’s instructions.

Orthologs of *C. elegans* genes involved in mitotic proliferation and cell cycle were identified in *B. malayi* genome (Bmal-4.0, Ghedin *et al*. 2007) using BLASTP and the open-access WormBase ParaSite website (Harris et al., 2009). Primer pairs were designed according to Primer3 version 0.4.0 (Untergasser et al., 2012) such as the forward and reverse primers were hybridized in two different exons to avoid background genomic DNA contamination (see Table S1 for details). All primers were commercially synthesized by Eurofins Genomics and their efficiency was close to 100%. Primers were diluted to a final concentration of 200 nM in the master mix. Amplifications were performed using the Brilliant III Ultra-Fast SYBR Green QPCR Master Mix (Agilent Technologies, #600882), Mx3000P instrument (Agilent Technologies), and MxPro QPCR Software (Agilent Technologies) using the option SYBR Green (with Dissociation Curve) according to manufacturer’s instructions. The PCR cycling program consists of a pre-amplification cycle of 3 minutes at 95°C followed by 40 amplification cycles of 20 seconds at 95°C then, 20 seconds at 60°C and a dissociation/melt cycle of 1 minute at 95°C followed by 30 seconds at 60°C and 30 seconds at 95°C. For each primer pairs, quantitative measurements were carried out in triplicate on three independent biological replicates. Among the two reference genes tested (Table S1), Bma-Anc-1 gene was the best one identified with NormFinder tool (Andersen et al., 2004) and was used to normalize gene expression.

#### Nucleotide quantification

Flash-frozen *B. malayi* and *A. vitae* females in liquid nitrogen were used to crush ~20 mg of worms per replicate (corresponding to 15 *B. malayi* females and 4 *A. viteae* females) on dry ice followed by an extraction with 2*1mL of methanol/water (80/20 v/v) at -40°C containing 240 μL of fully 13C-labeled cellular extract IDMS (used as internal standard). After centrifugation (5 min, 10000g, 4°C), the supernatants were recovered and evaporated to remove the solvents. Samples in triplicates were suspended in 240 μL of ultra-pure water prior to the mass spectrometry analysis. Samples were analyzed at the MetaToul facility (Metabolomics & Fluxomics Facitilies, Toulouse, France) by ion chromatography (ICS 5000+ system, Dionex, Sunnyvale, CA, USA) coupled with a 4000 QTrap triple quadrupole mass spectrometer (ABSciex, Framingham, MA, USA) equipped with a Turbo V source (ABSciex) for electrospray ionization (Kiefer et al., 2007). Intracellular metabolites were analyzed in the multiple reaction monitoring (MRM) mode, and the isotope dilution mass spectrometry (IDMS) method (Wu et al., 2005) was used to ensure accurate quantification. Fragmentation was done by collision-activated dissociation using nitrogen as the collision gas at medium pressure.

### Quantification and Statistical Analysis

All statistical analyses and graphics were carried out using R 3.1.3 (R Development Core Team 2010). Non-parametric Wilcoxon rank-sum tests were used for two samples comparisons, except for the relative gene expressions that were analysed with Student’s t-tests. Exact p-values and sample sizes are indicated in the corresponding figures.

## SUPPLEMENTAL ITEM TITLES AND LEGENDS

**Supplemental Table 1**

List of genes and primers used for qRT-PCR

**Supplemental Figure 1**

**The tetracycline treatment efficiently eliminates *Wolbachia*.**

Distal parts of ovaries dissected from wild-type and *Wolbachia-depleted* females, stained with DAPI -cyan- and propidium iodide -red-. The latter stains preferentially the *Wolbachia* DNA while the DAPI reveals the host DNA (i.e. in the wild-type ovary). The combination of these DNA dyes allows an easy look at the localization and the presence of the symbionts. Distal part to the left, the yellow arrows point to the DTCs. Scale bar = 5 μm.

**Supplemental Figure 2**

**Characterization of the *Brugia malayi* spermatogenesis in wild-type and *Wb*(*-*) males.**

Confocal images of (A) A dissected testis stained with DAPI. The yellow arrow points to the DTC, and the distal part is attached to the pharynx -green dotted lines-. (B) A distal part of a testis from a male incubated in EdU for 8 hrs corresponding to the area highlighted by green box in (A), and stained respectively with DAPI, an anti-phosphorylated histone H3 -PH3-antibody, EdU and followed by a merge of the Edu -yellow- and PH3 -magenta-channels. (C) Proliferative zones of linearized testes from wild-type and *Wolbachia-depleted* males, stained with DAPI and an anti-PH3 after an EdU incubation. Scale bars = 50 μm. (D) Left panels: Products of the second meiotic division from wild-type and *Wolbachia-depleted* males show proper segregation patterns. The actin (green) highlights the connecting residual body. Right panels: a DAPI stain on mature spermatocytes from wild-type (top) and *Wolbachia-depleted* (bottom) reveals the five, aligned chromosomes. Scale bars = 1 μm.

**Supplemental Figure 3**

**Scoring of apoptotic nuclei in the proliferative zone and apoptotic indexes.**

(A) Segment of a female proliferative zone stained with DAPI -red- and an anti phospho histone 3 -green-. The small round, DAPI-bright and PH3-negative nuclei are apoptotic - yellow arrows-. (B) Apoptotic indexes in the PZ of ovaries from *B. malayi* females collected from untreated or tetracycline-treated gerbil hosts. Antibiotic treatments were administered *per os* for 4 days.

**Supplemental Figure 4**

**Heat maps of mitotic nuclei and nuclear densities distributions along ovarian proliferative zones from wild-type and *Wolbachia-depleted B.malayi* females.**

Distribution of (A) mitotic nuclei and (B) total nuclear counts along female PZ, per segments of 50 μm starting from the distal tip of the ovaries. Each line represents the analysis of one ovary. For wild-type ovaries n= 21, and n= 29 for *Wolbachia-depleted* conditions.

**Supplemental Figure 5**

**Laser ablation of the Distal Tip Cell does not prevent germline proliferation.**

(A) Impact of the laser ablation on the DTC in living worms. Bright field acquisitions during the experiment: 1 Before ablation, the DTC is clearly visible at the tip of the ovary -yellow inset-. 2 The localized laser impact heats up the cell and creates a transient air bubble -black in the image-. 3 Few seconds later the DTC appears destroyed. (B) Confocal images of a pair of ovaries from a wild-type female, DTC-ablated and spared, stained with an anti-PH3 - magenta- and DAPI -grey- 24 hrs post-ablation. (C) Quantification of PH3+ nuclei in both types of ovaries. (D) A Phalloidin stain -grey- reveals the actin and confirms the absence of DTC 24hrs after laser ablation -DAPI in blue, propidium iodide in magenta-.

**Supplemental Figure 6**

**Quantitative PCRs on *B. malayi* orthologs of cell cycle regulators controlling the germline proliferation in *C. elegans*.**

The fold change was normalized using *Bma anc-1* to measure expression levels for *Bma cycline E, cdk2* and *cdc25*.

**Supplemental Figure 7**

**Germline stem cells loose their quiescence upon *Wolbachia* removal.**

(A) Quantification of PH3+ nuclei in the 1st 150 μm of spared control and DTC-ablated ovaries, 20 hrs post-ablation and after a 4 hr treatment with colchicine 1mM. (B) Distal ovaries from wild-type and *Wb*(*-*) worms collected after 72 hrs of *in vivo* EdU treatment in gerbils, EdU in cyan, DAPI in grey. (C) Number of EdU-positive nuclei counted in the most distal 0 to 50 μm and 0 to 150 μm-long segments of ovaries from wild-type and *Wolbachia-depleted* females.

